# Age-Related Delay in Visual and Auditory Evoked Responses is Mediated by White and Gray matter Differences

**DOI:** 10.1101/056440

**Authors:** D. Price, L. K. Tyler, R. Neto Henriques, K. Campbell, N. Williams, M. Treder, J. R. Taylor, CamCAN, R. N. A. Henson

**Author notes:** Corresponding Author: MRC Cognition & Brain Sciences Unit, 15 Chaucer Road, Cambridge, CB2 7EF, UK.

## Abstract

Slowing is a common feature of ageing, yet a direct relationship between neural slowing and brain atrophy is yet to be established in healthy humans. We combine magnetoencephalographic (MEG) measures of neural processing speed with magnetic resonance imaging (MRI) measures of white- and gray-matter in a large (N=617, 18-88yrs), population-derived cohort (www.cam-can.org) to investigate the relationship between age-related structural differences and VEF and AEF delay across two different tasks. Using a novel technique, we show that VEFs exhibit a constant delay, whereas AEPs exhibit delay that accumulates over time. Visual delay is mediated by white-matter microstructure in the optic radiation, presumably reflecting increased transmission time, whereas auditory delay is mediated by gray-matter differences in auditory cortex, presumably reflecting less efficient local processing. Our results demonstrate that age has dissociable effects on neural processing speed, and that these effects relate to different types of brain atrophy.

## Introduction

Age-related declines in cognitive abilities like fluid intelligence and working memory are well-documented, and a major burden for both older individuals and the societies they inhabit^1^. One potential common cause for these cognitive declines is a general slowing of information processing speed^2^. This slowing can be measured by behavioural responses in various tasks, which has been related to age-related atrophy of white-matter^3^^–^^5^ and gray-matter^6^^–^^10^. Animal studies have proposed at least two mechanisms of age-related slowing: demyelination, which results in longer axonal transmission times between neurons^11^'^12^, and changes in the neuron itself (such as increased hyperactivity), which results in reduced neural responsiveness^13^. These animal studies reinforce the importance of changes in both white- and gray-matter, and raise the possibility that these changes cause different types of neural slowing. Despite these links between white-matter, gray-matter and behavioural slowing in humans, and between physiological changes and neural processing delays in animals, there is currently no evidence from studies of healthy ageing in humans that directly links differences in brain structure with differences in neural processing speed. Such evidence would provide important mechanistic insights into the causes of age-related slowing of information processing, and hence cognitive decline. We provide such evidence by combining magnetoencephalography (MEG) and magnetic resonance imaging (MRI) from a large sample of 617 population-derived healthy adults, distributed uniformly from 18-88 years of age, recruited from the Cambridge Centre for Ageing and Neuroscience (CamCAN) (www.cam-can.org).

Age-related slowing of the neural response evoked by simple visual stimuli such as checkerboards has been observed using event-related potentials (ERPs) recorded with electroencephalography (EEC)^14^^–^^17^. Similar effects have also been found for auditory stimuli^18^^–^^21^, complex visual stimuli such as faces^22-26^, and olfactory stimuli^17^^,^^27^. Many studies have reported effects of age on early components of the ERP/ERF (e.g. within 200ms of stimulus onset)^15^^,^^17^^,^^28^^–^^30^, while others have reported age effects on later components (typically 200- 800ms) without a corresponding increase in latency in the early components^18^^,^^23^^,^^25^^,^^26^^,^^31^^–^^34^. We propose that these reflect two distinct types of delay: *constant* and *cumulative* delay. Constant delay affects all time-points equally, equivalent to a temporal shift of the whole evoked response (both early and late components). Cumulative delay, on the other hand, increases with post-stimulus time, and therefore is easier to detect for late than early components. Despite reports of cumulative delay in the literature^18^^,^^24^, there has been no systematic comparison of constant and cumulative delay, and it is possible that they have different neuronal causes. Research on senescent monkeys suggests that age related delay of the VEF has a cortical origin^13^, while a review comparing retinal electroretinogram ERG, and cortical evoked potentials suggests that delays in humans originate in the retinogeniculostriate pathway^28^. Similarly, the causes of delay of the auditory evoked response may include contributions from peripheral, central auditory system or cortical functional deficits^35^^–^^39^.

The study of age-related delays in ERPs/ERFs is further complicated by inconsistent findings in the degree of age-related delay of both early and late components (for review, see^17^^,^^25^^,^^26^^,^^28^). There are several likely sources of this inconsistency. Firstly, differences in information processing demands across experimental tasks (such as attending to faces versus ignoring auditory oddballs) mean that evoked responses are difficult to compare across studies. While clearly providing important information in their respective domains, complex tasks like face recognition are likely to recruit multiple cortical systems, and it is unclear how each system contributes to the spatially-integrated signal recorded by EEG/MEG. Secondly, age-related delays may differ according to the type of stimulus, for example visual versus auditory stimuli, because age could potentially have differential effects on brain regions specialised for different sensory modalities. Even when results are consistent across different tasks and modalities, they are rarely compared directly within the same group of participants. Thirdly, previous studies have tended to compare small groups of young versus old volunteers, rather than examine continuous differences across the adult lifespan, and tended to use volunteers who are self-selecting, rather than being representative of the population.

Yet another important source of inconsistency across studies is the method of measuring delay. One common measure is the latency of the peak (maximum) of an evoked component. This measure is very sensitive to noise however (being based on a small number of time points), so other techniques pool over several time points^23^, for example using the fractional area latency^40^, or the slope or intercept of functions fit to parts of the evoked response. FFere we use all (peristimulus) time-points to estimate delay, improving robustness and sensitivity, and enabling estimation of second-order delay characteristics like constant and cumulative delay, which are not readily available from single peak latencies, and difficult to quantify when there are multiple, potentially overlapping temporal components.

To address some of the other limitations in the literature, we directly compare the effects of age across two types of task (passive and active) and on two types of stimuli (visual and auditory). Furthermore, we apply our novel method for simultaneously estimating constant and cumulative delay to a larger and more representative (opt-out) sample than is typically tested, spanning the whole adult lifespan. Finally, we relate these estimates of neural delay, for the first time, to structural estimates of both gray-matter and white-matter on the same individuals.

More precisely, we used simple stimuli (visual checkerboards and pure tones) that are likely to activate only a few brain regions, and compare two tasks that differ in attentional demand: i) a “passive” viewing/listening session in which visual or auditory stimuli are presented separately, and do not require a response, versus ii) an “active” task in which the same stimuli are presented simultaneously, and require a motor response. We also use MEG, which has the same temporal resolution as the EEG, but has the advantage of higher spatial resolution, because magnetic fields are less spatially distorted by biological tissue than electrical fields^41^, thereby increasing our ability to separate brain sources (see also^26^). The measures of brain structure came from three types of MRI contrast: Tl-weighted, T2-weighted and diffusion-weighted. The T1 and T2 data are combined in order to optimise estimation of local gray-matter (GM) volume. Diffusion data are optimised for estimation of the mean kurtosis (MK) of the tissue’s water diffusion^42^^,^^43^, which is believed to offer a sensitive metric of age-related changes of white-matter microstructure, such as changes of cell membranes, organelles and the ratio of intra and extra-cellular water compartments^44^. Moreover, in contrast to standard diffusion tensor measures, diffusion kurtosis measures are robust to regions with a high concentration of crossing fibres^43^^,^^45^.

Our main experimental hypotheses were that age is positively correlated with constant and/or cumulative evoked response delay in auditory and visual conditions, and that this relationship between age and delay is mediated by differences in white-matter microstructure or gray-matter volume.

## Results

### PCA-Derived Event Related Fields

We start with data from the passive task, in which auditory tones and visual checkerboards were presented in separate trials. In order to reduce our dataset to a small set of meaningful components and improve the signal-to-noise ratio, principal component analysis (PCA) was performed on the trial-averaged event-related fields (ERFs) for each stimulus-type, after concatenating the data in the time dimension (see Methods). The first principal component explained 48% of the variance for auditory stimuli and 28% of the variance for visual stimuli, and entailed a spatial component for all participants (Figure 1a) plus a separate time-course for each participant (Figure 1c). The individual timecourses were then averaged to create a template ERF for later model fitting (Figure 1d). We used multiple sparse priors (MSP)^46^ to localise the spatial component, which revealed a single peak in bilateral primary auditory cortex for the auditory stimuli and in bilateral extrastriate cortex for the visual stimuli (Figure 1a-b). Importantly, the ERP-images in Figure 1c show how the ERFs are delayed as age increases, with indication of a cumulative shift for auditory stimuli and a constant shift for visual stimuli; a difference that we formally quantify below.

**Figure 1.**
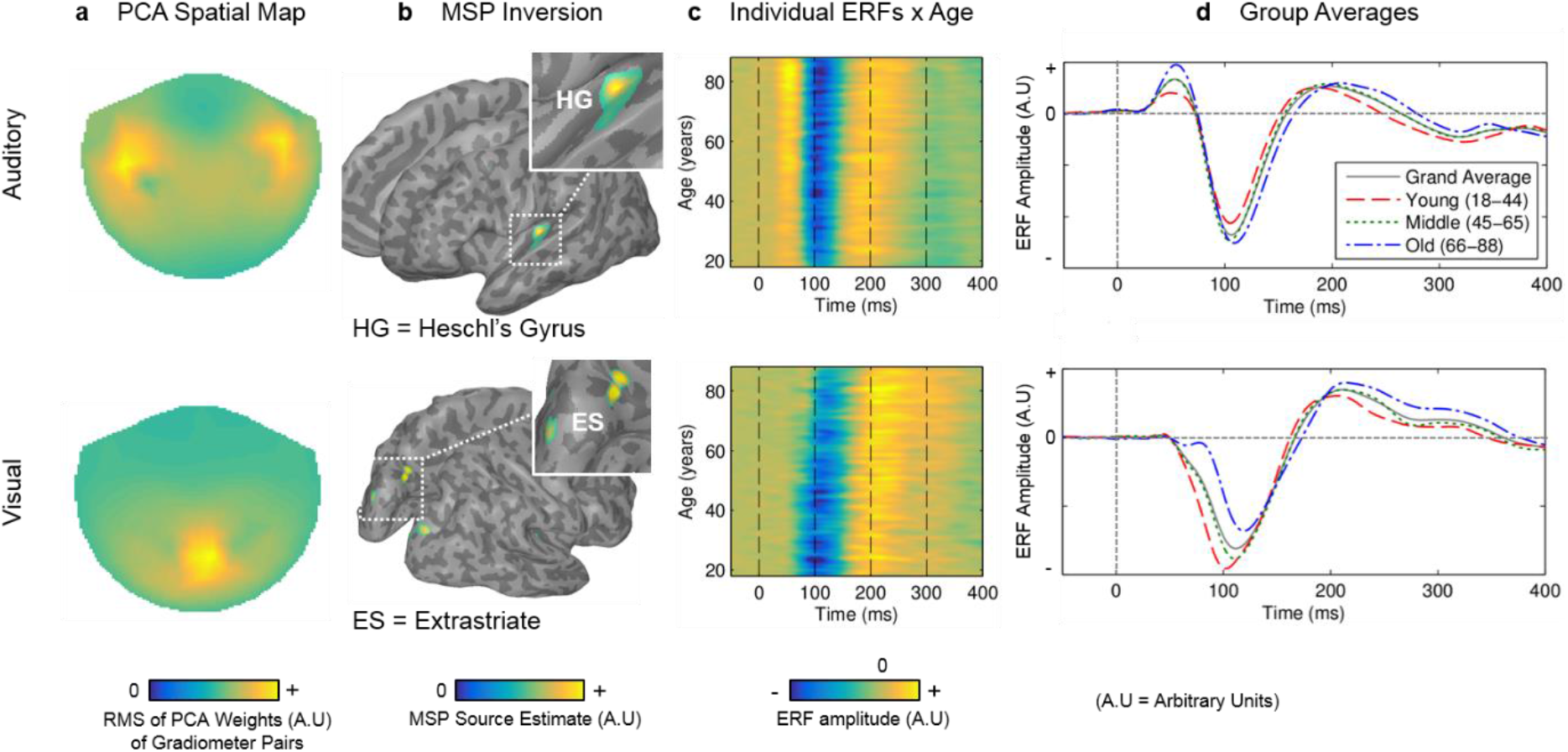
(a) 2D topographical MEG sensor plot of the first spatial component derived using PCA in the passive session. Values represent the root mean square (RMS) of each pair of gradiometers. (b) Group Multiple Sparse Prior (MSP) source reconstruction based on the spatial component shown in (a) (cluster peak MNI coordinates: right HG = [+38, −22, +8], left = [−38, −26, +8]; right V2 = [+14, −96, +20], left = [−14, −96, +20]; ES right = [+16, −74, +24], left = [−16, −74, +20]). (c) Heat-maps illustrating the time-courses for each participant from the first temporal component of the PCA. Data are smoothed in the y-direction for visualisation only (Gaussian width = 5 subjects), (d) Group average time-courses for each of three age groups (18-44, 45-65, 66-88yrs). Because all of our analyses are based on principal components, the y-axis of plots have arbitrary units.

In order to estimate constant and cumulative delay for each participant, a template fitting procedure was employed, in which the group average signal (gray line in Figure 1d) was fit to each participant’s ERF by a combination of temporal displacement (constant delay) and temporal linear dilation (cumulative delay). Using a local gradient ascent algorithm, these two parameters were adjusted until the linear model fit (R^2^) was maximised. Using R^2^ as the utility function simplifies the fitting procedure, and allows simultaneous estimation of the amplitude offset and amplitude scaling (see Methods).

Constant and cumulative delay estimates for the AEFs and VEFs were correlated with age, using robust regression after removing outlying values for each modality separately (Figure 2). In the visual condition, there was a significant effect of age on constant delay (R^2^=0.11, P<0.001, N=511), but no effect of age on cumulative delay (R^2^=0.00, P=0.884, N=511). In the auditory condition, on the other hand, there was a significant effect of age on cumulative delay (R^2^=0.14, P<0.001, N=567), but not on constant delay (R^2^=0.00, P=0.175, N=567). There was also a smaller but statistically significant increase in amplitude scaling of the AEF with age (R^2^=0.04, P<0.001, N=567), a small increase in auditory amplitude offset (R^2^=0.02, P=0.005, N=567), and an increase in visual amplitude offset (R^2^=0.09, P<0.001, N=501) (see supplementary Figure 6). For conversions of these results to ms/year, and comparison with traditional peak latency estimates, see supplementary Table 4.

**Figure 2.**
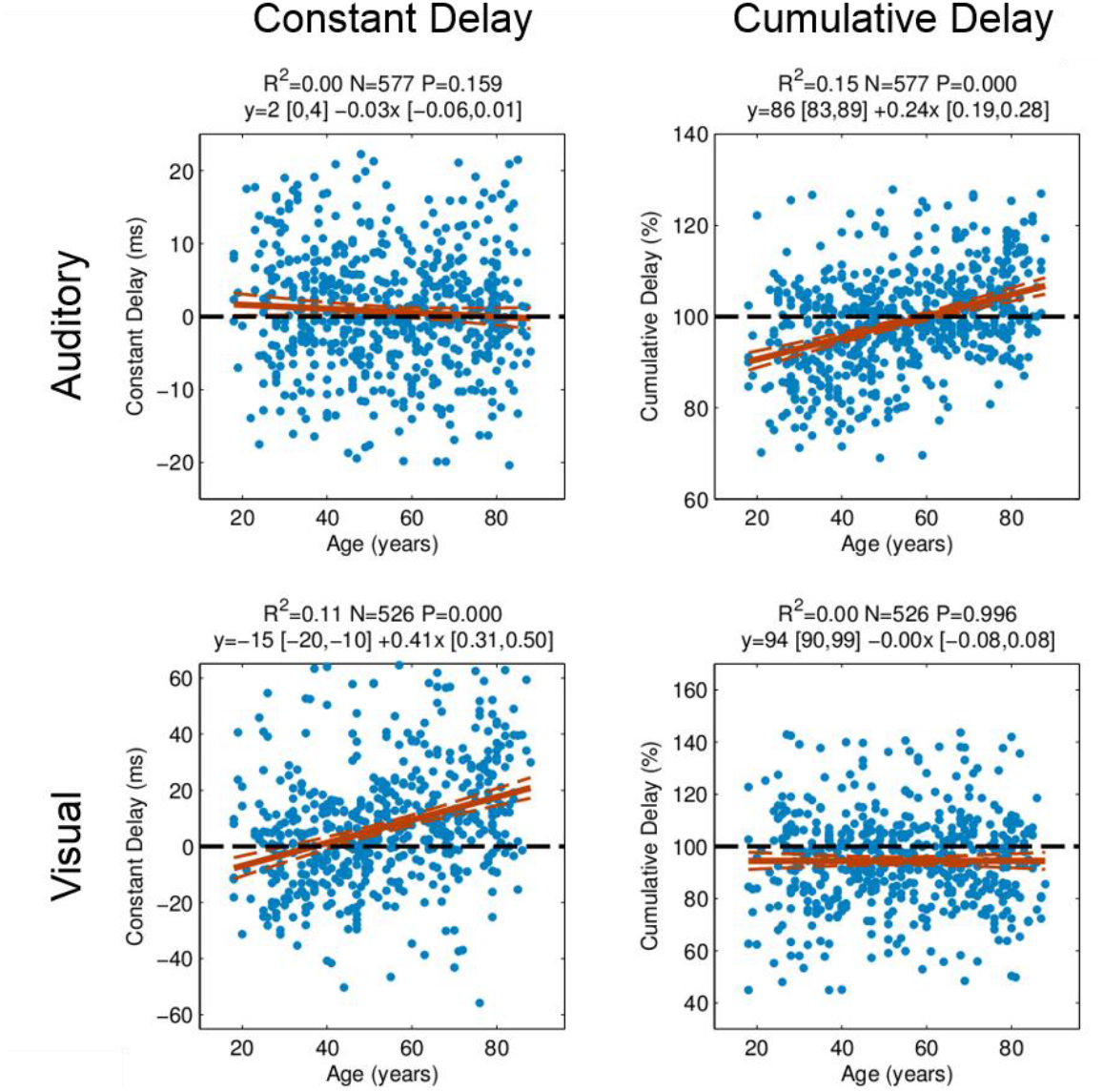
Robust regressions with age of the two delay parameters for each stimulus modality in the passive session. There is a significant effect of age on constant but not cumulative delay in the VEF, and a significant effect of age on cumulative but not constant delay in the AEF.

We also repeated the PCA and ERF fitting steps for the active task, in which the auditory and visual stimuli were presented simultaneously, and to which the participant responded with a key-press. This PCA produced similar results, though the simultaneous presentation of visual and auditory stimuli meant that the first spatial component reflected a mixture of responses to both stimulus-types (supplementary Figure 2). To better separate the auditory and visual responses in the active task, and aid comparison with the passive data, we applied the spatial weights of the PCA from the passive data to estimate the timecourses for each stimulus-type in the active task. Importantly, age had a significant effect on the visual constant delay (R^2^=0.03, P<0.001, N=473), but not visual cumulative delay (R^2^=0.00, P=0.959, N=473), and on the auditory cumulative delay (R^2^=0.09, P<0.001, N=513), but not auditory constant delay (R^2^=0.00, P<0.188, N=513), replicating the pattern of significant age effects in the passive session.

We also tested whether the task moderated the size of the age effect on the latency parameters. The effect of age on the auditory cumulative parameter was greater in the active task than the passive task, although no other delay parameters showed such an interaction between task and age (see supplementary Table 3). Nonetheless, for subsequent analyses below, we focus on the passive data, where the auditory and visual responses are more easily separated by PCA.

### Adjusting for sensory acuity, fit amplitude and fit error

It is possible that the above effects of age on neural delays are simply a consequence of the known age-related changes in sensory acuity (despite our screening criteria, and use of lenses to correct vision; see Methods). We therefore correlated the neural delay estimates against separate, standardised measures of auditory and visual thresholds (see Methods). Results are summarised in supplementary Table 1. There was a negative relationship between visual acuity and visual constant delay (r_s_=0.13, P=0.003, N=511), though this effect disappeared after adjusting for age (r_s_=−0.01, P=0.830, N=511). Similarly, auditory acuity was negatively related to auditory cumulative delay (r_s_=0.22, P=0.002, N=567), but not after adjusting for age (r_s_=−0.06, P=0.l49, N=567). Importantly however, both visual and auditory age-related neural delays remained significant after controlling for visual (r_s_=0.30, P<0.001, N=511) and auditory (r_s_=0.31, P<0.001, N=567) sensory acuity, respectively.

To check that the effects of age on neural delay estimates were not biased by effects of age on the estimated response amplitude (offset or scaling) or the estimated fit quality, we also repeated the partial correlation of neural delays with age after adjusting for these estimates. For both visual and auditory measures, the effects of amplitude scaling on delay were uncorrelated after controlling for age (auditory cumulative r_s_=.31, P<0.001, 582; visual constant r_s_=−0.05, P=0.245, N=511) and the effects of age on delay remained significant after controlling for amplitude (auditory cumulative r_s_=0.37, P=0.000, N=582; visual constant r_s_=0.34, P=0.000, N=511). The same was true for amplitude offset and fit error (see supplementary Table 1). Thus, there is no evidence that our age-related neural delays were caused by age-related differences in sensory acuity, response amplitude or offset, or goodness of template fit.

In summary, it appears that age has a qualitatively different effect on neural delay in auditory and visual modalities. Indeed, the auditory cumulative delay was only weakly correlated with visual constant delay (R^2^=0.02, P=0.005, N=491), and this effect vanished after controlling for age (R^2^=0.00, P=0.947, N=491), suggesting these two delay parameters are not intrinsically related, and are therefore likely to have separate underlying causes. To investigate possible causes, we turned to the MRI data on each participant.

### Mediation of Neural Latency by Structural Brain Measures

In order to test the hypothesis that brain structural changes account for some of the shared variance between age and neural delay, whole-head voxel-wise robust mediation analyses were performed^47^. Mediation analysis tests whether the relation – path c – between a predictor variable (X, age) and an outcome variable (Y, ERF delay) is significantly attenuated when the relation between X and a mediator variable (M, white matter microstructure or gray-matter volume) – path a – and the relation between M and Y – path b – are added to the model. Four separate models were tested at each voxel: one for each type of delay (auditory cumulative or visual constant) as the outcome and one for each brain measure (white or gray-matter) as the mediator. All models included total intracranial volume (TIV) as a covariate of no interest. Mediation effect sizes were computed for every voxel, and a voxel-wise false detection rate (FDR) of 5% applied in order to correct for multiple comparisons. This threshold was further Bonferroni-corrected for multiple comparisons across the four models. Finally, voxels were also required to i) show significant relations between age and mediator (path a) and between mediator and outcome (path b) at the same level of FDR correction, and ii) fall within gray-matter or white-matter masks.

The mediation effects of white-matter microstructure on the relationship between age and visual constant delay are displayed in Figure 3a. Two clusters of significant voxels were found in the left retrolenticular part of the internal capsule (RIC) (label 1), and the left posterior thalamic radiation (PTR) (label 2). These paths together form the optic radiation projecting from the lateral geniculate nucleus (LGN) to the primary visual cortex (VI). The cluster centred on left RIC extended to left superior corona radiata and left superior longitudinal fasciculus, although the mediation effects here were generally lower. The cluster visible in left PTR also extended to splenium of corpus callosum, connecting left and right occipital cortices. There was no evidence that white-matter mediated the effects of age on auditory cumulative delay.

**Figure 3:**
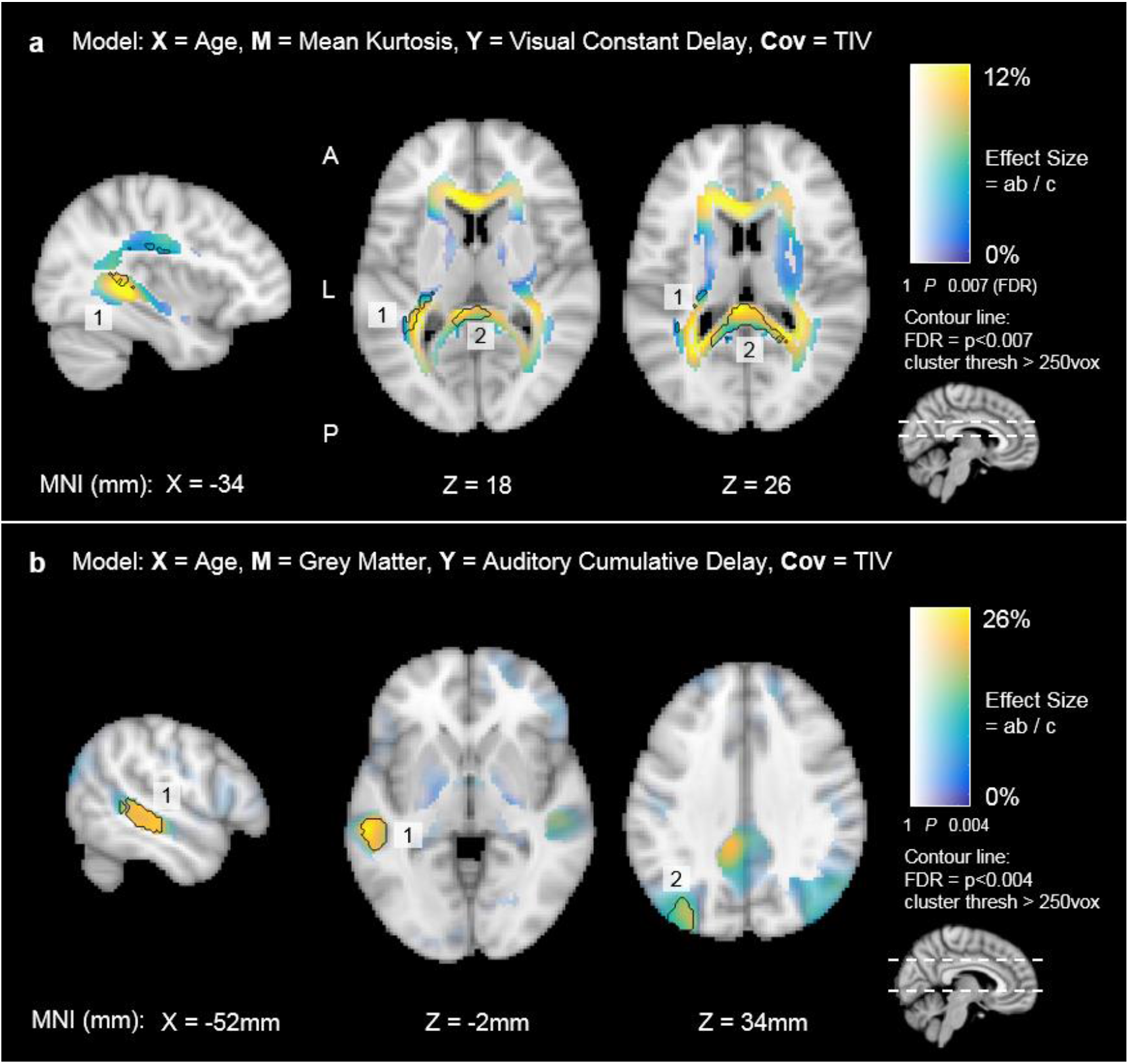
Whole-brain voxel-wise mediation analysis results. Voxel hue corresponds to the effect size (% mediation effect), while opacity corresponds to the univariate p-value. Clusters of at least 250 voxels that survive false detection rate (FDR) correction are marked with a black border, a) WM structure (mean kurtosis, MK) in the optic radiation (connecting LGN to VI) [label 1], and in splenium of corpus callosum [label 2], mediates the age versus visual constant delay relationship, b) GM volume in the left posterior superior temporal gyrus (STG) mediates the age versus auditory cumulative delay relationship [label 1], Another cluster was observed in the left superior lateral occipital cortex, a region not involved in auditory processing [label 2]. All effects indicate positive mediation (i.e., the age vs. delay relationship is attenuated when including the mediator).

Mediation effects of gray-matter on the relationship between age and auditory cumulative delay are displayed in Figure 3b. One cluster was found comprising the left posterior superior temporal gyrus (STG) (label 1), extending to middle temporal gyrus (MTG). Another cluster was observed in superior lateral occipital cortex (label 2), although effect sizes here were lower (peak = 12%) than in STG (peak = 26%). There was no evidence that gray-matter mediated the effects of age on visual constant delay.

## Discussion

Using a novel analysis technique on MEG data from a large, lifespan cohort, we discovered two distinct types of age-related neuronal delays in sensory evoked responses: a constant delay in the visual evoked magnetic field (VEF) to checkerboards and a cumulative delay in the auditory evoked magnetic field (AEF) to tones. These delays occurred regardless of whether participants were passively encountering auditory and visual stimuli separately, or actively responding to them when both were presented concurrently, demonstrating that these age effects occur under variable levels of attention. After controlling for common age effects, these two types of delay were uncorrelated across individuals, suggesting dissociable causes. In support of this, we found that white-matter microstructure (mean kurtosis, MK, from diffusion-weighted MRI), primarily in the optic radiation (LGN to VI), mediated the effect of age on the visual constant delay, whereas gray-matter volume (as estimated from T1- and T2-weighted MRIs), primarily in the posterior superior temporal gyrus (STG), mediated the effect of age on the auditory cumulative delay. We discuss these findings in light of prior behavioural, neuroimaging and animal studies.

### Visual Constant Delay

The age-related neural slowing in the VEF and its relationship to white-matter microstructure in the optic radiation point to a delay in the arrival time of information communicated from the LGN to VI. Several aspects of our results support this conclusion. Firstly, the main generator of the VEF was located in prestriate (V2) and extrastriate visual cortex. Secondly, the slowing was characterised by a constant delay of the entire ERF, so that both early (∼50-1 50ms) and late (∼150-400ms) components of the evoked response were affected, consistent with a delayed arrival time. Thirdly, the whole-brain voxel-wise mediation analysis revealed that microstructural differences in the left retrolenticular internal capsule (RIC) and posterior thalamic radiation (PTR) partially account for the age-related delay. The RIC in particular is a major junction of fibres that transmit information from the LGN to visual cortex (and our use of diffusion kurtosis likely increased our sensitivity to regions with such crossing or fanning fibres). Furthermore, the lack of cumulative delay suggests that, once the information reaches visual cortex, it is processed at a normal rate (cf. auditory results below).

These results contrast with findings in the senescent macaque monkey^13^, which revealed no white-matter atrophy, and no delay in the LGN-V1 pathway. Instead, spike-timing delays were related to increased neuronal excitability and an increase in transmission time from VI to V2. One reason for this discrepancy may be differences in the neural measures (e.g., spiking rates versus the local field potentials recorded with MEG); another may be that humans have different neurobiological ageing profiles to macaques, highlighting the importance of investigating ageing *in-vivo* in humans. Our results do not preclude the possibility that other information-carrying fibres and thalamic nuclei play a role in visual processing delays, but clearly point to age-related differences in white-matter microstructure being at least partially responsible for delays in information processing associated with healthy human ageing. Further, our findings support a review comparing pattern-evoked electroretinogram (ERG) and cortical evoked potentials that concluded age-related visual deficits are the result of disruption to the retinogeniculostriate pathway^28^.

### Auditory Cumulative Delay

The age-related cumulative delay observed in the auditory evoked response, along with the lack of constant delay, points to a different mechanism than that associated with the visual response. We suggest that this cumulative delay reflects a deficit in local processing within auditory cortex, specifically recurrent interactions between primary auditory cortex and higher order auditory regions. We again base this claim on several aspects of our analysis. Firstly, the main source of the AEF was the primary auditory cortex, in-line with evidence that this region responds to pure tones^48^. This delay was mediated by gray-matter volume in a higher order auditory region, namely the superior temporal gyrus (STG). We also observed a cluster in left superior lateral occipital cortex, but since this region is not involved in processing of auditory stimuli, we suggest that collinear decline between this region and STG is responsible for this result, rather than any true functional link. The STG is too distant from the primary source to be considered the generator of the AEF, but is certainly anatomically connected to primary auditory cortex^49^, suggesting that interactions with STG are responsible for the temporal dilation of the AEP. Indeed, the auditory system is organised into a functional hierarchy, beginning with processing of simple sounds (tone and frequency) in primary auditory cortex, before further processing in surrounding “belt” regions in the superior temporal lobe^48^^,^^50^^,^^51^. It is difficult to say with certainty, but the cumulative delay observed in the present study may therefore be linked to a disruption of neural function (possibly neural recovery times^52^) and dynamic interactions between primary auditory cortex and higher processing regions, resulting in delay that worsens over the duration of the evoked response, but does not affect the arrival time of information from the peripheral auditory system. Finally, we cannot completely rule out the contribution of central auditory processing deficits on our results^19^^,^^53^. However, central deficits are linked with age-related amplitude increases of the AEP, while our study demonstrated an independence of amplitude and latency. Further research is clearly needed to determine the contribution of central auditory processing deficits in cumulative delay of the AEP.

This idea of delays in local cortical interactions may also explain recent studies^22^^,^^23^ that have reported age-related cumulative delay for information accrual during processing of simple vs. complex visual stimuli after approximately 90-120ms. These findings cannot be explained by the visual constant delay found here, because one would expect simple and complex stimuli to be equally delayed, leaving no net delay when contrasting them. We suggest that increasing stimulus complexity results in a greater dependence on recurrent communication between brain regions responsible for face discrimination. The speed at which this occurs, and therefore the temporal profile of the evoked response, would then be dependent on the integrity of those neural networks. Future studies directly comparing the delay profiles of responses across simple and complex discrimination tasks may help to shed light on these effects. Furthermore, computational modelling would help to understand the different neural mechanisms that could result in constant versus cumulative delay in different sensory systems.

### Other factors affecting age-related delay

One potential additional cause of age-related delay in evoked responses is the well-known age-related change in sensory acuity. If sensory acuity were an adequate predictor of neural delay, we would expect a relationship between acuity and delay to remain even after adjusting for age. However, while auditory acuity was weakly correlated with auditory cumulative delay, when accounting for age, this correlation disappeared. The same was true for the relationship between visual acuity and visual constant delay. Most importantly, the correlations between age and both visual and auditory delays remained significant after adjusting for sensory acuity. These results suggest that sensory acuity does not play a significant role in our findings, which is consistent with previous studies arguing that optical and retinal factors cannot fully account for age-related delays in the visual evoked response^14^^–^^16^^,^^22^^,^^28^.

### Assumptions

The template fitting method described here is conditional on several assumptions about the generators of the ERF. Firstly, in applying PCA to our entire dataset we assume spatial stationarity of the evoked responses across the age range. It is possible that age related changes in structural morphology result in changes in spatial distribution of the evoked fields. However, it is unlikely that any such changes would have such striking effects on the timecourse of the evoked response. Furthermore, we compared young and old participants and did not observe any evidence of spatial non-stationarity (supplementary Figure lb), nor did we find any qualitatively different results when using a template derived from just young or just old participants (supplementary Figure 7). Secondly, we assume temporal stationarity of all delay parameters. That is to say that the equations derived during fitting apply to the entire timeseries (epoch), rather than there being distinct, short-lived evoked components within that epoch that are subject to different age-related delays. This rests on the hypothesis that evoked responses in MEG/EEG are the result of sustained, dynamic interactions between neuronal sub-populations within and across brain regions^51^^,^^54^, producing damped oscillations, rather than being distinct and transient evoked “components” at the peaks and troughs of ERF/ERP waveforms. If this assumption is true, then changes to the physical characteristics of those neuronal populations (such as GM and WM integrity) are likely to lead to temporally extended delay characteristics. Indeed, we performed extensive simulations (supplementary Figure 3) to demonstrate that, when temporal stationarity holds, our template method is much more sensitive (in presence of noise) than more traditional peak-based, peak-to-peak or fractional area measures of latency. We also confirmed that our conclusions hold when performing those more traditional peak-based analyses on separate time-windows (supplementary Figure 4). Finally, we investigated whether there was evidence in our data against temporal stationarity. To do this, we repeated our template fitting approach on a shorter time window from 0-140ms that only covered the first visual peak and first two auditory peaks (excluding the later, more dispersed components). The results show that largely the same pattern of age-effects on delay is observed in this early window as it is in the entire epoch (see supplementary Figure la), supporting the present assumptions.

We found evidence that the effect of age on some latency estimates does depend on the task in which latency is estimated, in that the slope of the relationship between auditory cumulative delay and age was higher in the active task than passive task. We cannot tell from our two paradigms whether this effect of task reflects 1) whether or not a response to the visual/auditory stimuli is required, and/or 2) whether or not the visual and auditory stimuli are simultaneous. This is direct evidence for the concern raised in the Introduction that divergent results in the literature may reflect the use of quite different tasks. Nonetheless, the dissociable effects of age on visual constant versus auditory cumulative delay held across both of the present tasks (the slope for auditory cumulative delay was simply greater in the active task), suggesting that the basic age effects on visual and auditory delay are somewhat invariant to the presence of the motor component of the task, or the concurrent presentation of auditory and visual stimuli.

### Caveats

While our results go beyond previous studies in testing for age-related delays across both visual and auditory modalities and across two different tasks, our findings are nonetheless restricted to simple visual and auditory stimuli, which may not generalise to more complex stimuli that require more extensive neural processing. This might explain the age-related cumulative delay found previously in the ERP to faces^23^, which likely involves a greater degree of recurrent processing between multiple visual cortical regions responsible for face perception.

Secondly, caution is warranted over interpretation of our mediation analysis. Mediation analysis is a statistical approach that cannot properly determine causality in the same way that an intervention might (e.g., to lesion parts of the optic radiation and test effects on visual constant delay). Furthermore, mediation results from cross-sectional studies cannot be interpreted solely in terms of the ageing process^55^, and have alternative explanations such as cohort effects. Whatever the precise role of age, our findings nonetheless demonstrate that there are at least two types of neural processing delay, which are unrelated across individuals, and have different relationships with white- and gray-matter in the brain structures associated with that processing.

Our study is the first to investigate the relationship between delays in human evoked responses and brain structure (as measured by MRI). As such, we had no specific hypotheses about the relative roles of grey-versus white-matter, or different brain regions. Thus, even though our whole-brain search revealed statistically significant and mechanistically interpretable results, our findings should be regarded as exploratory. Future studies could take a more confirmatory approach to testing the role of specific structural properties in sensory processing delays.

Finally, note that we are not claiming that our present white- and gray-matter findings are a complete account of age-related slowing. Our voxel mediation effect sizes explained at most 26% of the age-related variance in response delays. It is possible that a larger proportion of variance in neural delay could be explained by combining brain measures across voxels, but even then, full mediation would be surprising since several other factors not measured here, such as altered neurotransmitter concentrations^56^ and central auditory processing deficits^53^ are also potential contributory factors. Nonetheless, our findings represent an important step forward in demonstrating dissociable types of age-related neuronal slowing, and generating mechanistic hypotheses for their causes.

### Conclusion

In summary, the present work fills a missing gap in the literature: evidence that in healthy humans, age-related delay of the electrophysiological response to stimulation is due to structural differences of functionally-relevant brain regions responsible for the transfer and processing of information. We have taken this a step further to show that neural delay should not be thought of as a unitary concept that affects all brain regions equally. Instead, ageing appears to be associated with regionally-specific changes in characteristic neural responses, which are likely due to heterogeneous age-related changes across anatomical structures.

## Methods

### Participants

Participants were recruited from a healthy population-derived sample from the Cam-CAN study (www.cam-can.org; see Shafto et al.^57^ for a comprehensive explanation of the study design and experimental protocol). Ethical approval for the study was obtained from the Cambridgeshire Research ethics committee. Prior to the home interview, individuals give written informed consent for the study. Written informed consent is also given by participants at each scanning session. Participants were excluded based on several criteria: Mini Mental State Examination (MMSE) < 25; failing to hear a 35dB lkH tone in either ear; poor English language skills (non-native or non-bilingual speakers); self-reported substance abuse and serious health conditions (e.g. major psychiatric conditions, or a history of stroke or heart conditions); or MRI or MEG contraindications (e.g. ferromagnetic metallic implants, pacemakers, or recent surgery). Participants that did not take part in both the MEG and MRI sessions were also excluded. The final sample of N=617 had an age range of 18-88 years at the time of first contact. For a post hoc analysis of power (number of participants needed to detect the present effect sizes), see supplementary Figure 5. In addition to screening tests at home interview stage, participants took both a visual (Snellen sight test, with vision correction) and auditory acuity test immediately preceding the MEG scan (see Shafto et al.^57^ p7 for details). The scores from these tests were also used in later statistical analysis of age-related neural delay to control for the possible confounds of age-related differences of visual/auditory acuity (8 participants with missing acuity data were removed) (see supplementary Table 1).

### Audio / Visual Tasks

The visual stimulus consisted of two circular checkerboards presented simultaneously to the left and right of a central fixation cross (34ms duration × 60 presentations). The diameter of each circular checkerboard subtended an angle of 3°, their centroids were separated by 6° on the horizontal plane, and the checks had a spatial frequency modulation of 2 cycles per degree. Visual stimuli were presented using a Panasonic PT-D7700 DLP projector (1024×768, 60Hz refresh rate) outside of the MSR, projected though a waveguide onto a back-projection screen placed 129cm from the participant’s head. The stimulus onset was adjusted for the 34ms (2 refreshes at 60FFz) delay induced by the projector. The auditory stimulus was a binaural tone (300ms duration; 20 presentations of 300Hz, 600Hz, and 1200Hz; 60 total presentations), with a rise and fall time of 26ms, and presented at 75dB sound pressure level (measured using an artificial ear). The stimulus onset was adjusted for the 13ms delay for the sound to reach the ears. The first session involved the active task, in which participants were presented with both types of stimuli concurrently, and asked to respond by pressing a button with their right index finger after stimulus onset. The reaction times (RTs) are analysed in supplementary Table 2, though note that the active task was not explicitly speeded. The stimulus-onset asynchrony varied randomly between 2s and 26s, to match an fMRI version of same task (see Taylor et al.^58^ for more details). The session lasted 8min and 40s in total.

The second session was the passive task. Participants experienced separate trials with either the visual or auditory stimulus, which they were asked to passively observe (no response required). Visual and auditory trials were pseudo-randomly ordered, with an SOA that varied randomly between 0.8s to 2s. Given the simultaneous presentation, evoked responses were adjusted by the average delay of 23ms (13ms auditory and 34ms visual). The session lasted approximately 2 minutes.

### PCA Based Latency Analysis

The 2D (time × sensor) matrices for each participant were concatenated along the time dimension. PCA was performed with columns as signals and rows as observations to produce a set of time domain signals for each trial-type. The *n*^th^ principal component was then reshaped to give individual trial averages for each participant. The principal component weights represent the degree to which each channel contributes towards the *n*^th^ principal component. Given that the relationship is linear, the weights can also be used for source localisation. Since the simple sensory-evoked sources are likely to be distributed across multiple regions, we used multiple sparse priors (MSP), implemented in SPM12, which is capable of recovering multiple sparsely distributed generators of ERFs^46^. Each participant’s MRI was coregistered to their MEG data using three anatomical fiducial points (nasion, and left and right pre-auricular points) that were digitised for the MEG data and identified manually on the MRIs. Lead fields were calculated using a singleshell model based on the deformed spheres approach^59^.

### ERF Fitting

The aim of the fitting procedure was to obtain estimates for two types of delay: constant delay, defined as delay that affects all time points equally (modelled using a 0^th^ order delay parameter); and cumulative delay, defined as delay that is linearly dependent on the time point at which it is measured (modelled using a 1^st^ order delay parameter). Note that the delay terms could be of any order, but limiting the delay estimates to 0^th^ and 1^st^ order terms reduces the danger of over-fitting. ERF fitting was performed using in-house code written in MATLAB. Firstly, a template ERF was computed from the data as the trial-averaged ERF for a given principal component, averaged across all participants. Because this template represents the group average, approximately equal numbers of participants will have negative and positive delay parameters when fitting their individual ERFs. A template fitting algorithm was designed to iteratively alter the temporal characteristics of the template signal ***s***(***t***) by:

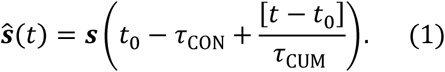

where *τ_CON_* is the constant delay, *τ_CUM_* is the cumulative delay and *t*_0_ is a stationary point in time for any value of *τ_cum_*, and was constant across all tests (see below). Cubic spline interpolation was used to obtain *ŝ* for any given set of delay parameters.

#### Fitting Procedure

For each individual time-course ***S*(*t*)**, the parameters were adjusted iteratively using a local maximum gradient ascent algorithm until the convergence criteria was satisfied. Starting parameters were chosen to correspond to the null hypothesis that no delay would be observed compared to the group average (*τ_CON_* = 0; *τ_CUM_* = 1) and delay parameters were adjusted by a predefined quantity (*τ_CON_* = +/−20*ms*; *τ_CUM_* = +/−10%), giving 4 fits per iteration. A value of *t*_0_ = 50*ms* was fixed on the basis of the typical latency for information to reach sensory cortices (e.g. Plm). Fit was determined using linear regression, and the parameter values that gave the best *R*^2^ fit were chosen as starting points in the next iteration. Using regression simplified the fitting procedure, since amplitude scaling and mean were determined from the regression model’s beta estimate and intercept, respectively. If none of the parameters resulted in a better fit than the current best fit, then the magnitude of the parameter adjustments were reduced by a factor of 0.75 and the process was repeated. Convergence was achieved when the *R^2^* fit improvement over the current best fit was less than le-6. This method helped to reduce the occurrence of spurious outliers, because the gradient ascent was constrained to converge into the maxima closest to the starting point. Results were visually inspected to ensure optimal fitting was achieved. Outlier parameter values were handled during statistical testing, described below.

### Robust Regression

For each delay parameter, outliers were identified based on the boxplot rule (+/−1.5 times the inter-quartile range, IQR) and removed from the analysis. An outlier in either the constant or cumulative delay estimate resulted in the participant being rejected from the analysis for a given experimental condition (visual or auditory). For all correlation analyses, robust methods were employed to control for remaining extreme values. Robust regression was implemented using MATLAB Statistical Toolbox (LinearModel.fit). A bisquare weighting function with a tuning constant of 4.685 was used to weight cases based on their residual error from an ordinary least squares fit, then repeated until convergence.

For the purpose of comparing our results with those in the literature using more conventional measures, the linear equation describing the age-related change in constant and cumulative delay can be converted back to peak-latency, *l*, by combining the age vs. constant/cumulative delay regression equations (from Results) with Equation 1 (where *t* becomes the peak-of-interest, *α* is age, and *τ_CON_* and *τ_CUM_* are replaced with the linear equations obtained from our main analysis):

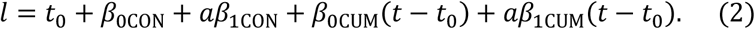

Separating the constant and age dependent terms gives the coefficients of the linear equation *l = β*_0_ + *αβ*_1_:

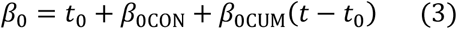

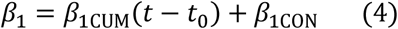

where *t* is the peak-of-interest (e.g. for P2m, *t* = 200 - obtained from the ERF template), and *β*_0*con*_, *β*_0*cum*_, *β*_1*con*_ and *β*_1cum_ are the intercepts and slope betas from the age vs. delay regression equations. Confidence intervals can also be converted using the same formula.

### MRI Data

MRI data were acquired using a Siemens 3T TIM TRIO (Siemens, Erlangen, Germany) with a 32-channel head coil at the MRC Cognition & Brain Sciences Unit (CBU), Cambridge, UK. Anatomical images were acquired with a resolution of 1mm^3^ isotropic using a T1-weighted MPRAGE sequence (TR: 2250ms; TE: 2.98ms; TI: 900ms; 190Hz; flip angle: 9°; FOV: 256 × 240 × 192mm; GRAPPA acceleration factor: 2; acquisition time of 4 minutes and 32 seconds), and a 1mm^3^ isotropic T2-weighted SPACE sequence (TR: 2800ms; TE: 408ms; flip angle: 9°; FOV: 256 × 256 × 192mm; GRAPPA acceleration factor: 2; acquisition time of 4 minutes and 32 seconds). Diffusion-Weighted Images (DWIs) were acquired with a twice-refocused spin-echo sequence, with 30 diffusion gradient directions for each of two b-values: 1000 and 2000 s/mm^2^, plus three images acquired with a b-value of 0. These parameters are optimised for estimation of the diffusion kurtosis tensor and associated scalar metrics. Other DWI parameters were: TR=9100 milliseconds, TE=104 milliseconds, voxel size = 2mm isotropic, FOV = 192mm × 192mm, 66 axial slices, number of averages=l; acquisition time of 10 minutes and 2 seconds.

All MRI data were analysed using the SPM12 software (www.fil.ion.ucl.ac.uk/spm), implemented in the AA 4.2 batching software (https://github.com/rhodricusack/automaticanalysis: for complete description of data and pipelines, see Taylor et al.^58^).

The T1 and T2 images were initially coregistered to the MNI template using a rigid-body transformation, and then combined in order to segment the brain into 6 tissue classes: GM, WM, cerebrospinal fluid (CSF), bone, soft tissue, and residual noise. The GM images were then submitted to diffeomorphic registration (DARTEL) to create group template images, which was then affine-transformed to the MNI template. To accommodate changes in volume from these transformations, the GM images were modulated by the Jacobean of the deformations to produce estimates in MNI space of the original local GM volume.

The DWI data were first coregistered with the Tl image and then skull-stripped using the BET utility in FSL (http://fsl.fmrib.ox.ac.uk/fsl/fslwiki/). Linear fitting of a higher-order tensor was then used to estimate mean kurtosis (using in-house code). Images of the diffusion metrics were then normalised to MNI space using the DARTEL+affine transformations from the T1+T2 pipeline above.

### MEG Scanning and Pre-processing

Data were collected continuously using a whole-head Elekta Neuromag Vector View 306 channel MEG system (102 magnetometers and 204 planar gradiometers) (Elekta, Neuromag, Helsinki, Finland), located in a light magnetically shielded room (MSR) at the CBU. Data were sampled at 1kHz with a highpass filter of 0.03Hz. Recordings were taken in the seated position. Head position within the MEG helmet was estimated continuously using four Head-Position Indicator (HPI) coils to allow for offline correction of head motion. Two pairs of bipolar electrodes were used to record vertical and horizontal electrooculogram (VEOG, HEOG) signals to monitor blinks and eye-movements, and one pair of bipolar electrodes to record the electrocardiogram (ECG) signal to monitor pulse-related artefacts. Instructions and visual stimuli were projected onto a screen through an aperture in the front wall of the MSR. Participants were given MEG-compatible glasses to correct their vision. Auditory stimuli were presented binaurally via etymotic tubes. Motor responses were made via a custom-built button box with fibre optic leads.

**Figure 4:**
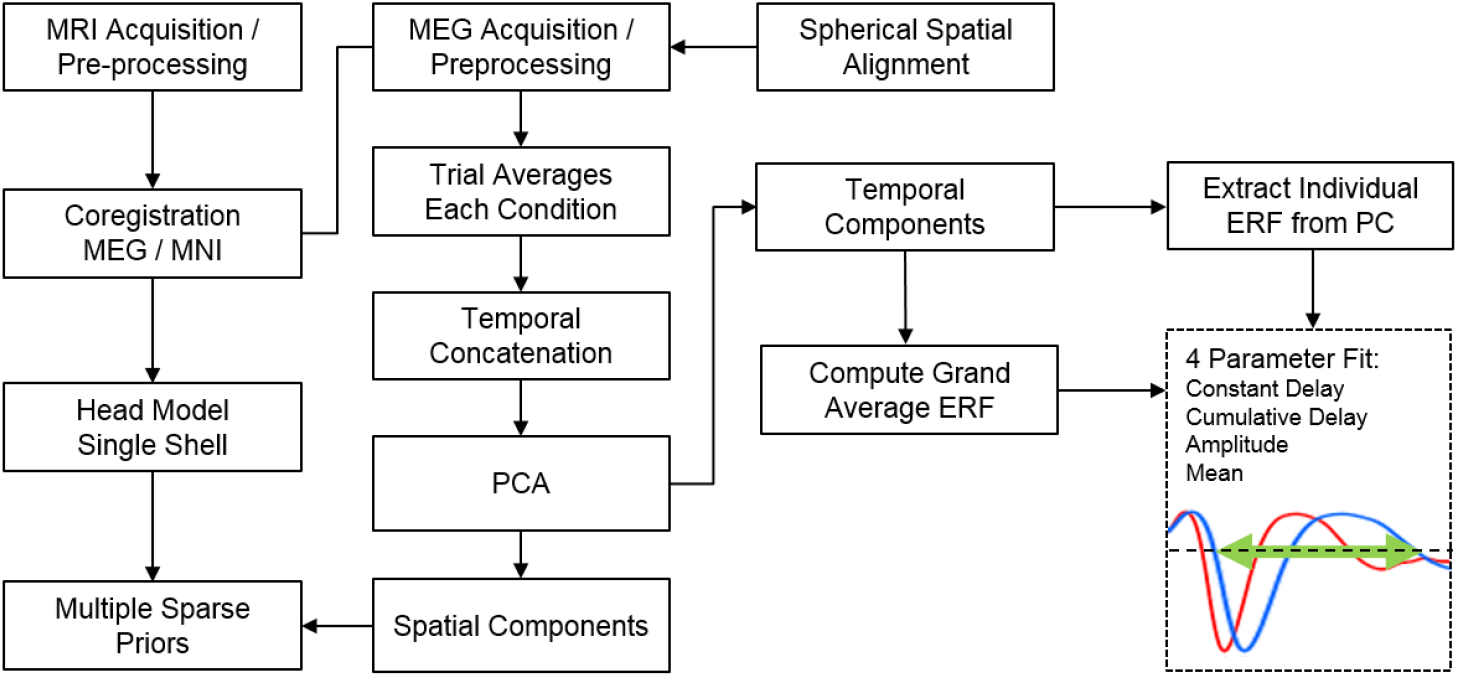
Flow chart diagram illustrating the processing steps involved in the analysis of MEG data.

Temporal signal space separation (tSSS; MaxFilter 2.2, Elekta Neuromag Oy, Helsinki, Finland) was applied to the continuous MEG data to remove noise from external sources and from HPI coils (correlation threshold 0.98, 10-sec sliding window), for continuous head-motion correction (in 200-ms time windows), and to virtually transform data to a common head position (‘-trans default’ option with origin adjusted to the optimal device origin, [0, +13, −6]). MaxFilter was also used to remove mains-frequency noise (50Hz notch filter) and to automatically detect and virtually reconstruct any noisy channels. Data were then imported into Matlab using SPM12. We identified physiological artefacts from blinks, eye-movements, and cardiac pulse using the logistic Infomax independent components analysis (ICA) algorithm implemented in EEGLAB. This was done by identifying those ICs whose time courses and spatial topographies correlated highly with reference time courses (correlation greater than three standard deviations from mean) and spatial topographies (correlation greater than two standard deviations from mean), respectively, for each of the above artefact types (run via in-house code, “detect_ICA_artefacts.m” available fromhttp://www.mrc-cbu.cam.ac. uk/people/rik.henson/personal/analysis/).

Data were then filtered using a two-pass 5^th^ order Butterworth filter (l-32Hz) implemented in fieldtrip, epoched (time window: −100-500ms peristimulus time), and the average baseline (-100-0ms) was subtracted from the data. Trial averaged responses of evoked amplitude were computed for each channel, participant and condition of the experiment. This resulted in a 2-dimensional matrix for each participant (channels x time). All subsequent analyses were carried out using data from the 204 gradiometer channels, since these are more sensitive to superficial sources than the magnetometer sensors, and the sensory cortices of interest here are superficial. ERF fitting on both the passive and task sessions was performed on the principal components of the channel level data (see *PCA based latency analysis* below).

### Whole-Brain Voxel-Wise Mediation Analysis

Images in MNI space of Mean Kurtosis (MK) from the DWI pipeline, and of local gray-matter volume (GMV) from the T1 and T2 pipeline, were smoothed (12mm FWHM) and entered into a whole-brain voxel-wise robust mediation analysis implemented using the M3 Mediation Toolbox (http://wagerlab.colorado.edu)^47^. A three path model was used with age as the predictor variable (X), delay as the dependent variable (Y) (outliers removed as above), and anatomical data as the mediator variable (M). Since total intracranial volume (TIY) was correlated with age (R^2^ = 0.01, P=0.03), it was also included as a covariate to control for the possible confounding effects of head size on structural statistics.

Mass univariate robust mediation was computed per voxel using the “robust fit” option of the M3 toolbox (10,000 bootstrapped samples per voxel) to generate path data (paths: a = X−>M, b = M−>Y, c’ = X−>Y, c = ab+c’), and associated p-values calculated per voxel. Four models were tested, all with age as the predictor (X) and TIV as covariate of no interest, which correlated with age (R^2^=0.01, P=0.03, N=617). The mediator variable (M) was voxel data from either the white or gray-matter images. White-matter volumes were masked using the JHU ROIs (all ROIs were included); gray-matter volumes were masked using the Harvard Oxford gray-matter atlas. Only those voxels falling within the mask were entered into the mediation model. The false detection rate (FDR) threshold for each image was calculated from the resulting p-value maps using the M3 mediation toolbox. Statistical maps were thresholded according to these p-value thresholds, and clusters with fewer than 250 suprathreshold voxels were excluded. Mediation effect sizes were calculated using the formula, *M_effect_* = 100*ab/c*, which represents the mediation effect on the c path of including M in the model (resulting in c’) as a percentage of the total direct effect (c). A value of 100% indicates full mediation.

### Data Availability

Data from the CamCAN project is available from the managed-access portal at http://camcan-archive.mrc-cbu.cam.ac.uk. subject to conditions (see website) Analysis scripts are available in supplementary materials accompanying this publication.

## Acknowledgements

The Cambridge Centre for Ageing and Neuroscience (Cam-CAN) research was supported by the Biotechnology and Biological Sciences Research Council (BB/H008217/1); R.N.H. is additionally supported by the UK Medical Research Council (MC_A060_5PR10). We are grateful to the Cam-CAN respondents and their primary care teams in Cambridge for their participation in this study. We also thank colleagues at the MRC Cognition and Brain Sciences Unit MEG and MRI facilities for their assistance. Finally, we thank Dr Rousselet and two other reviewers for their helpful comments.

**Conflict of Interest:** None

## Contributions

CamCAN conceived the study. CamCAN piloted the study and collected the data. J.R.T., N.W., D.P., and R.H. collated and preprocessed the data. D.P. wrote the in-house MEG analysis software and analysed the data. D.P., R.N.H., L.K.T., N.H., K.C., N.W., M.T., J.R.T. wrote the paper.

## Footnotes

*Cam-CAN corporate author list:* The Cam-CAN corporate author consists of the project principal personnel: Lorraine K Tyler, Carol Brayne^5^ Edward T Bullmore^5^ Andrew C Calder^5^ Rhodri Cusack^5^ Tim Dalgleish^5^ John Duncan^5^ Richard N Henson^5^ Fiona E Matthews^5^ William D Marslen-Wilson^5^ James B Rowe^5^ Meredith A Shafto^5^’ Research Associates: Karen Campbell^5^ Teresa Cheung^5^ Simon Davis^5^ Linda Geerligs^5^ Rogier Kievit^5^ Anna McCarrey^5^ Abdur Mustafa^5^ Darren Price^5^ David Samu^5^ Jason R Taylor^5^ Matthias Treder^5^ Kamen A Tsvetanov^5^ Janna van Belle^5^ Nitin Williams’^5^ Research Assistants: Lauren Bates^5^ Tina Emery^5^ Sharon Erzinglioglu^5^ Andrew Gadie^5^ Sofia Gerbase^5^ Stanimira Georgieva^5^ Claire Hanley^5^ Beth Parkin^5^ David Troy^5^’ Affiliated Personnel: Tibor Auer^5^ Marta Correia^5^ Lu Gao^5^ Emma Green^5^ Rafael Henriques^5^’ Research Interviewers: Jodie Allen^5^ Gillian Amery^5^ Liana Amunts^5^ Anne Barcroft^5^ Amanda Castle^5^ Cheryl Dias^5^ Jonathan Dowrick^5^ Melissa Fair^5^ Hayley Fisher^5^ Anna Goulding^5^ Adarsh Grewal^5^ Geoff Hale^5^ Andrew Hilton^5^ Frances Johnson^5^ Patricia Johnston^5^ Thea Kavanagh-Williamson^5^ Magdalena Kwasniewska^5^ Alison McMinn^5^ Kim Norman^5^ Jessica Penrose^5^ Fiona Roby^5^ Diane Rowland^5^ John Sargeant^5^ Maggie Squire^5^ Beth Stevens^5^ Aldabra Stoddart^5^ Cheryl Stone^5^ Tracy Thompson^5^ Ozlem Yazlik^5^’ and administrative staff: Dan Barnes^5^ Marie Dixon^5^ Jaya Hillman^5^ Joanne Mitchell^5^ Laura Villis^5^.

